# Research Waste in ME/CFS

**DOI:** 10.1101/133926

**Authors:** Sonia Lee

## Abstract

**Objective:** To compare the prevalence of selective reporting in ME/CFS research areas: psychosocial versus cellular.

**Method:** A bias appraisal was conducted on three trials (1x psychosocial and 2x cellular) to compare risk of bias in study design, selection and measurement. The primary outcome compared evidence and justifications in resolving biases by proportions (%) and ORs (Odds Ratio); the secondary outcome determined the proportion (in %) of ME/CFS grants at risk of bias.

**Results:** NS (cellular study) was twice as likely to present evidence in resolving biases over PACE (psychosocial trial) (OR = 2.16; 65.6% vs 46. 9%), but this difference was not significant (*p* = 0.13). However, NS was five times more likely to justify biases over PACE (OR = 4.76; 46.9% vs 15. 6%) and this difference was significant (*p* = 0.0095; *p <* 0.05). PACE was weak in place (operational aspects 32%) and NS for data practices (37%). The proportion of grants were more biased in PACE (72%) than NS (28%) for evidence, and also more biased in PACE (86%) than NS (14%) for justifications.

**Conclusion:** Psychosocial trials on ME/CFS are more likely to engage in selective reporting indicative of research waste than cellular trials. Improvements to place may help reduce these biases, whereas cellular trials may benefit from adopting more translatable data methods. However, these findings are based on two trials. Further risk of bias appraisals are needed to determine the number of trials required to make robust these findings.

## 1. Background

Research waste in clinical trials are seen in outcomes that are not published, or in selective reporting of incidental and spurious findings that cannot be reproduced or translated in practice. When outcomes are not published: resources are wasted, research is stilted, and the study protocol cannot be validated nor repudiated in future protocols. Reviews on publication rates indicate: 50% of randomised trials are not published (Kasenda *et. al.*, 2014); 88% for cohort studies (Bogert *et al.*, 2015); and 50% for pre-clinical and clinical studies (Schmucker *et al.*, 2014). On the other hand, selective reporting is suspected when data is fabricated (intentionally misrepresented); or falsified (intentionally manipulated) in favour of a desired outcome. The potential causes of selective reporting include: poor recruitment, irrelevant endpoints, biased selection criteria and discontinuation, for instance: of 1017 RCTs, 25% were discontinued, and of those, 9.9% were discontinued due to poor recruitment (Kasenda *et. al.*, 2014). When outcomes are not published, authors are contacted for missing data in instances of imputing data in unpublished trials (systematic review). However in selective reporting, even if the reported outcomes appear distinctly remarkable: *p* hacking (extremely good *p* values); file drawer problem (only positive results), it is difficult to substantiate who is responsible for it, and whether it was intentional; and whether institutional enquiries into research misconduct are worth pursuing if proven to be futile against existing policies, and run the risk of polarising research communities.

As clinical trials become more complex, there is increasing concern selective reporting is harder to detect, and unforeseen complexities may escalate between the oversight bodies that monitor research integrity (eg. issues of research misconduct) versus the autonomy which allow research communities to freely conduct their own research. This review seeks to demonstrate these complexities in Chronic Fatigue Syndrome (CFS).

## 2. ME/CFS

Myalgic Encephalomyelitis (ME) and CFS are not yet considered distinct diagnoses, but have been in the past (White *et. al.*, 2007). The time to onset and the causes of these fatigue-like symptoms are confined to case studies (low evidence), and is still debated among experts. Nevertheless, both are diagnosed when there is an absence of fatigue-related disorders, and the patient achieves a minimum threshold score for ME/CFS in at least one fatigue questionnaire eg. The Chalder Fatigue Scale; The Krupp Fatigue Severity Scale; DePaul Symptom Questionnaire etc. (Board on the Health Select Populations, 2015).

In the UK, Cognitive Behavioural Therapy (CBT) and Graded Exercise Therapy (GET) are proposed treatment regimens for ME/CFS to reduce the symptoms of fatigue (NICE guidelines), and are based on the results of a randomised trial (PACE: Pacing, graded Activity, and Cognitive behaviour therapy) on ME/CFS patients (*n* = 641) conducted between March 2005 and November 2008. It recommends 12 to 15 sessions of Cognitive Behavioural Therapy (CBT; Fatigue: *n* = 161, *p* = 0.0136, *p <* 0.05) over 52 weeks; or 12 to 14 sessions of Graded Exercise Therapy (GET; Fatigue: n = 159, *p* = 0.0013, *p <* 0.05) over 52 weeks (White *et. al.*, 2011). ME/CFS support groups have rejected this treatment regimen due to harms from post-exertion malaise after GET, and no improvements after CBT. Biomolecular findings further support these claims with evidence of cellular level harms detected after GET (Cook *et. al.*, 2017), and have proposed biomarkers that are unique to ME/CFS (Fluge *et. al.*, 2016). The consensus is that ME/CFS is a complex, multi-faceted disorder that requires a multi-disciplinary approach, and aetiologies at multiple angles, such as: gut microbiota, hormonal, endocrine and immune functions. However, psychosocial angles can also offer important insights when studies are designed based on evidence. The following are lessons learnt from PACE, and recommendations on evidence-based study designs that may facilitate biomolecular studies, and potentially salvage psychosocial perspectives from branching off into research waste.

## 3. Methods

### 3.1. Search strategy

Search terms “myalgic encephalomyelitis”, “cognitive”, “behaviour”, “graded exercised therapy”, “adaptive pacing therapy”, “gene”, “cell”, “clinicaltrials.gov” were automatically mined from PubMed using E-utilities on a UNIX platform with no date restriction (fig. 1: sample code). The number of articles, authors and grants were tabulated by year in Excel. All articles, authors, and grants were included, and none were excluded. The search strategy collated MeSH terms for two research trends: Psychosocial versus Cellular. Articles were not scoped (included or excluded) for quality to observe only for research trends, and to minimise selection bias in future studies that may choose to replicate this search strategy. This search strategy did not scope for treatment effects as done in systematic reviews, but on research trends in selecting high impact trials for a bias appriasal.

**Figure 1:**
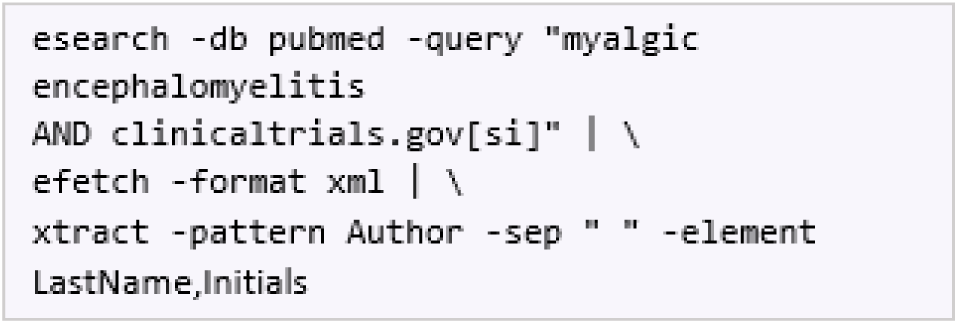
Sample search code in E-utilities (PubMed).

### 3.2. Bias appraisal

The author conducted a bias appraisal (an extended version of Cochrane’s bias appraisal tool) on three articles based on search outcomes: year and impact factor (fig. 2): 1. PACE: psychosocial interventions (GET, CBT, and Adaptive Pacing Therapy) to represent psychosocial trends (White *et. al.*, 2011); 2. A neural study (NS) on post-exertion malaise after GET to represent cellular trends (Cook *et. al.*, 2017). 3. A gut study (GS) on profiling gut microbial differences in ME/CFS individuals (Giloteaux *et. al.*, 2017) to also represent cellular trends. In table 1, biases were cate-gorised by: “study design”, “selection” and “measurement.” Each potential bias was rated by the author with a plus (+) or a minus sign (-) to indicate whether a study presented evidence (E) or a justification (J) for resolving a potential bias. The first two columns E and J rated PACE (White *et. al.*, 2011). The next two columns E and J rated GS (Giloteaux *et. al.*, 2017); followed by ratings for NS (Cook *et. al.*, 2017). The column “Neural Study” offered an example of each rating from Cook and colleagues’ (2017) paper. The column “Potential Biases” defined these biases in public health terms. The far right column with the letters “T”, “P”, “D”, “R”: Theory (theories and models used in the trial); Place (operational conduct); Recruitment (participant recruitment); Data (data practices) were collated to predict the areas of strengths and weaknesses (selective reporting) in each trial.

**Figure 2:**
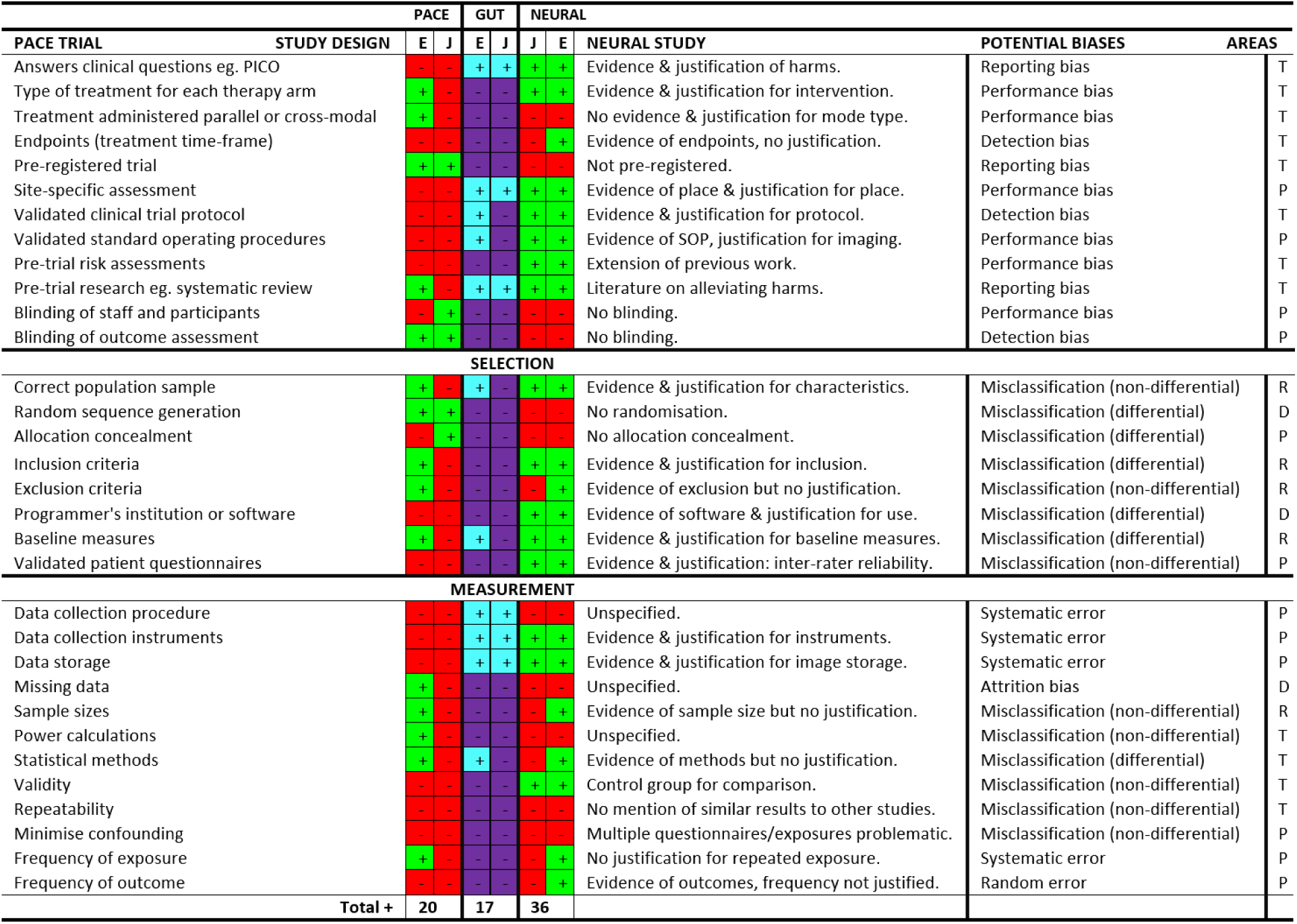
Summary of bias ratings. “+” sign indicates evidence or justifications present in the trial, “-” sign indicates it was not present. Areas “T”, “P”, “D”, “R” stand for Theory, Place, Data and Recruitment to represent the potential weak areas in the trial.

### 3.3. Statistical analysis

1. Primary outcome ME/CFS research trends: psychosocial or cellular were compared using a 2x2 contingency table to determine the strength of evidence (table 1) and justifications (table 2). Columns E and J from the bias appraisal (fig. 2) were tallied and imputed into two contingency tables, and its proportions were compared in deriving the Odds Ratio (OR). The OR determined the strength of evidence or justifications in resolving biases between the two research trends. It also determined the likelihood (in%) of evidence or justifications that were present in each research trend. Finally, *Z* tests (two-tailed) were performed to assess whether the use of evidence or justifications were significantly different between the two research trends.
2. Secondary outcome The OR (primary outcome) was applied to the total number of ME/CFS grants (search strategy) declared in each research trend (psychosocial versus cellular) to compare the proportion (in %) of grants at risk of bias.
3. Software All data were imputed and analysed by the author using a web-based clinical trials calculator (Centre for Clinical Research and Biostatistics, The Chinese University of Hong Kong) and verified for accuracy in another web-based, effect-size calculator (Campbell Collaboration).

**Table 1:**
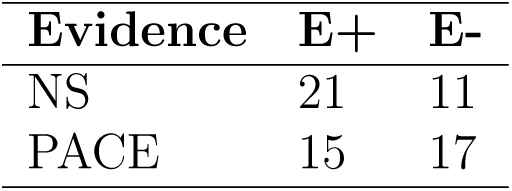
2x2 contingency table comparing evidence tallied from figure 2 (bias appraisal). NS stands for Neural Study; E+ for total plus ratings; E-for total minus ratings.

**Table 2.**
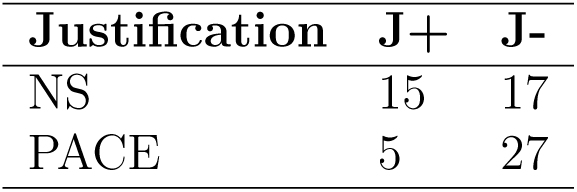
2x2 contingency table comparing justification tallied from figure 2 (bias appraisal). NS stands for Neural Study; J+ for total plus ratings; J-for total minus ratings.

| **Justification** | **J+** | **J-** |
| --- | --- | --- |
| NS | 15 | 17 |
| PACE | 5 | 27 |

## 4. Results

The search strategy identified 1750 published articles for psychosocial ME/CFS research and 1015 for cellular ME/CFS research between the dates 1951 to 25 March 2017 (day the search was performed). All articles were included in observing research trends: psychosocial versus cellular (fig. 1). Of interest were altmetric scores scoped manually in a psychosocial trial from 2011 (PACE; White *et. al.*, 2011); a cellular study from 2016 (NS; Cook *et. al.*, 2017); and another cellular study from 2016 (GS; Giloteaux *et. al.*, 2016). All articles were selected for popularity (altmetric scores) to represent ME/CFS research trends: 1. Psychosocial: PACE trial (White *et. al.*, 2011); 2. Cellular: Neural study (Cook *et. al.*, 2017); 3. Cellular: Gut study (Giloteaux *et. al.*, 2016) however, did not undergo any further analysis, since pre-trial conditions (eg. serum samples) from the bias appraisal (table 1) were significantly different in scope to the other two studies *ie.* it did not conform to an *intention to treat* experimental design.

### 4.1. Risk of bias appraisal

1. Study design The study design for PACE was unclear, it was neither a randomised controlled trial (no control group) nor a cohort study (randomised); whereas NS had a control group and was likely a case-control study. All of which may have influenced performance biases, attrition biases and other operational biases towards or away from the null. PACE had four treatment arms all ending in 52 weeks but these endpoints were not justified. NS also did not justify measuring neural endpoints in 3 days. Hence both studies were potentially biased on whether treatment effects were measured at an optimal level for a true effect. PACE registered the trial (ISRCTN54285094), NS did not, however, it did specify the site in which the intervention took place, whereas PACE did not, which are potential causes of attrition and reporting biases particularly when trial results can not be verified. NS based the intervention off prior work and used objective instruments (imaging scans) for measuring outcomes of an appropriate sample size; PACE on the otherhand, recruited a large sample size for an early phase trial, with no evidence of pre-trial risk assessments on standard operating procedures and clinical trial protocols which normally recruit small sample sizes in early phase trials (so to limit risks on the population of interest). The study design for GS was unclear, it did not conform to an intention to treat experimental design. It had a control group which were matched to cases, but it did not specify the number of specimens, patients and controls recruited by the physician, and if diagnostic tests *ie.* blood draws were done after an already confirmed ME/CFS diagnosis, which are all potential causes of performance, attrition and other biases.
2. Selection bias NS presented evidence and justifications for population sampling, baseline measures between cases and controls (patient characteristics), and validated questionnaires prior to conducting the study, but it did not conceal treatment allocations despite including a control group. This is highly problematic and can lead to selection biases consistent in case studies without a control group (low evidence). Conversely, if NS used a case-control design, it fell short of blinding and a good coverage of cases obtained from all facilities (not just one) to minimise selection bias. PACE presented better clinical protocols but did not justify population sampling, sample size, selection criteria, baseline measures, questionnaires and software. GS presented very little evidence and justifications in selection eg. it did not specify the number of participants and baseline measures that were relevant to sourcing serum samples. All of which are potential causes of selection bias.
3. Measurement bias All three studies were scant on evidence and justifications for data collection, analysis and dissemination. PACE did not specify who collected data, the types of software used, or whether the interventions were curative (given there were no treatment endpoints) and relevant to clinical practice. NS did not specify on which specialists assessed the imaging scans and whether the imaging scans were sufficiently sensitive and specific in diagnosing ME/CFS. Also, there was no blinding (if case-control), neither did it specify the number of participants screened per questionnaire (eg. information about lost controls) which often leads to biases in underestimating the prevalence of exposure in controls, and overestimate the Odds Ratio in favour of cases (those with ME/CFS). GS presented evidence and justifications for assessing data in a controlled lab environment, but it did not specify any missing data or contaminated samples, any lab errors, confounding or overlapping of RNA sequences with other disease conditions. It also used a rank statistical method, an unspecified machine learning method, a sub-sampling method for validating an unspecified model, all of which were unclear and potential causes of measurement bias.

### 4.2. Evidence ratings

NS was twice as likely to be supported by evidence compared to PACE (OR = 2.16). NS strength of evidence was at 65.6% and 33% relative to justifications; for PACE 46.9% and 23% relative to justifications (fig. 4). However, there was no significant difference between the two studies in presenting evidence to resolve biases (*p* = 0.13; *p >* 0.05).

**Figure 3:**
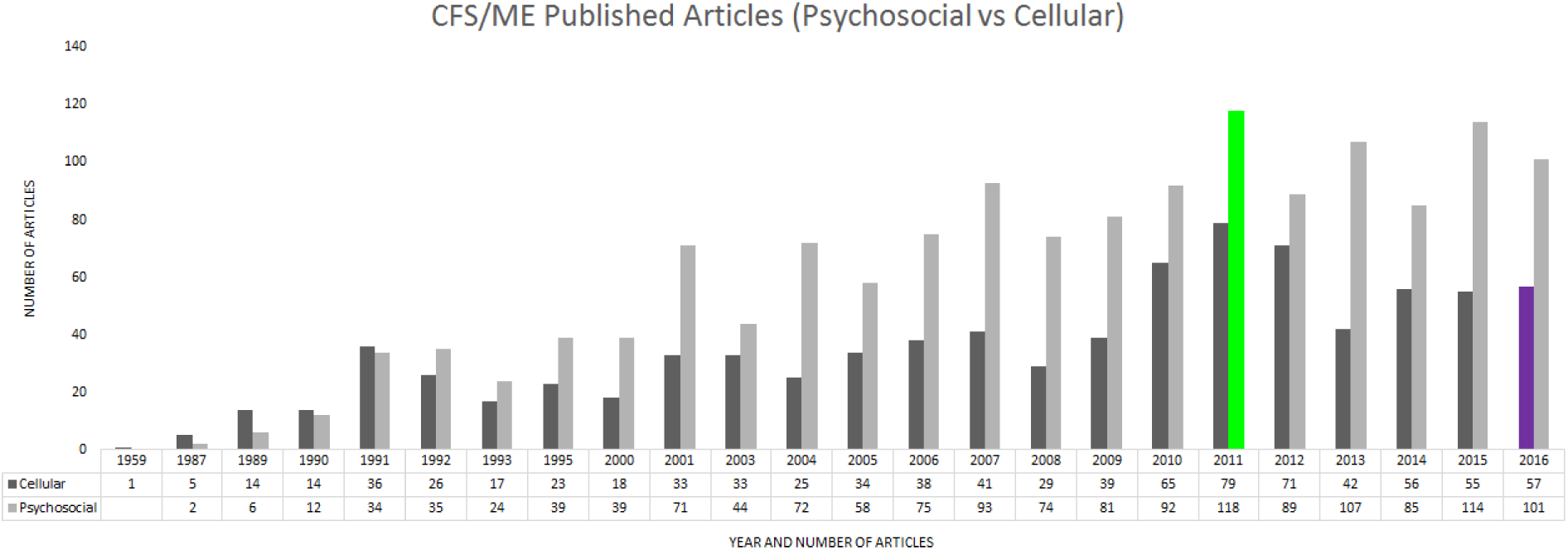
Search results comparing research trends: psychosocial versus cellular by year and number of published articles. The highlighted portions denote which year studies were selected for the bias appraisal.

**Figure 4:**
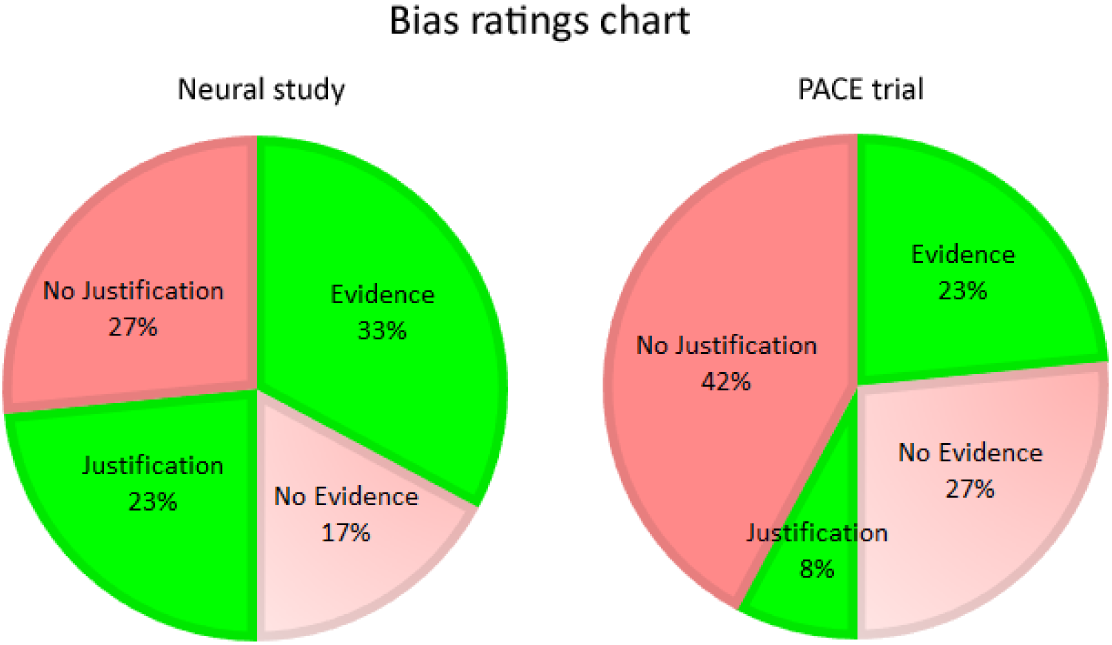
The proportion of bias in evidence and justifications in NS and PACE.

### 4.3. Justification ratings

NS was five times more likely to address biases compared to PACE (OR = 4.76). Justifications present in NS were at 46.9% and 23% relative to evidence; for PACE 15.6% and 8% relative to evidence (fig. 4). Unlike evidence, there was a significant difference in presenting justifications to resolve biases in favour of NS (*p* = 0.0095; *p <* 0.05).

### 4.4. Proportion of bias

Measurement bias was the most prevalent of all biases: PACE (39%) and NS (46%); followed by bias in study design: PACE (33%) and NS (29%); followed by selection bias: PACE (28%) and NS (25%) (fig. 5).

**Figure 5:**
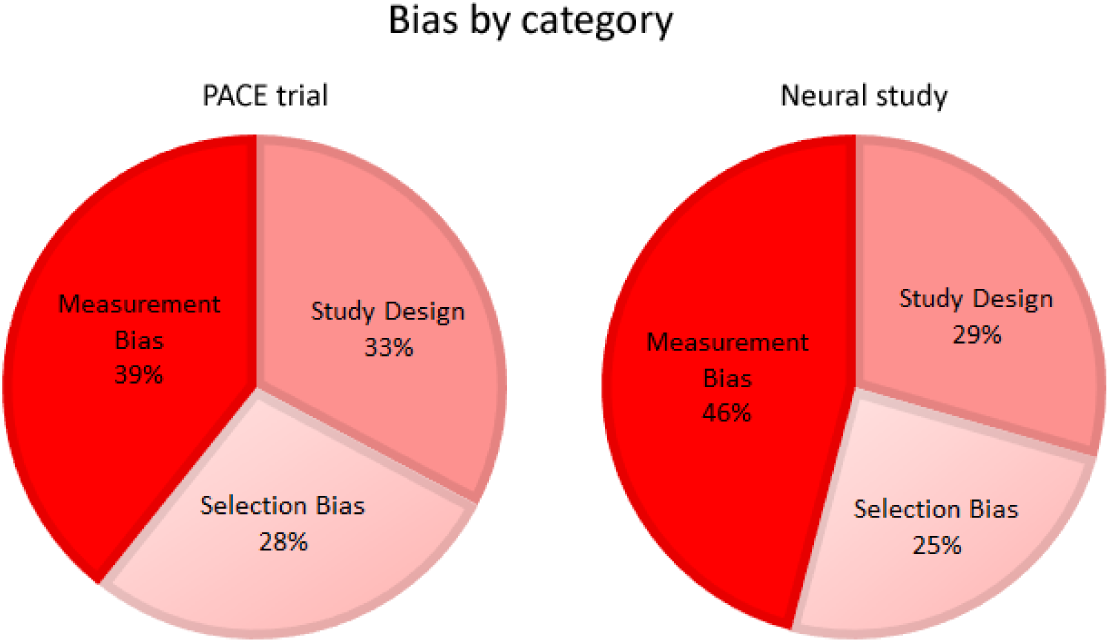
The proportion of bias by category in PACE and NS.

### 4.5. Study weaknesses

PACE was weak in place (32%) and theory (28%), whereas NS was weak in data (37%) and place (28%) (fig. 6).

**Figure 6:**
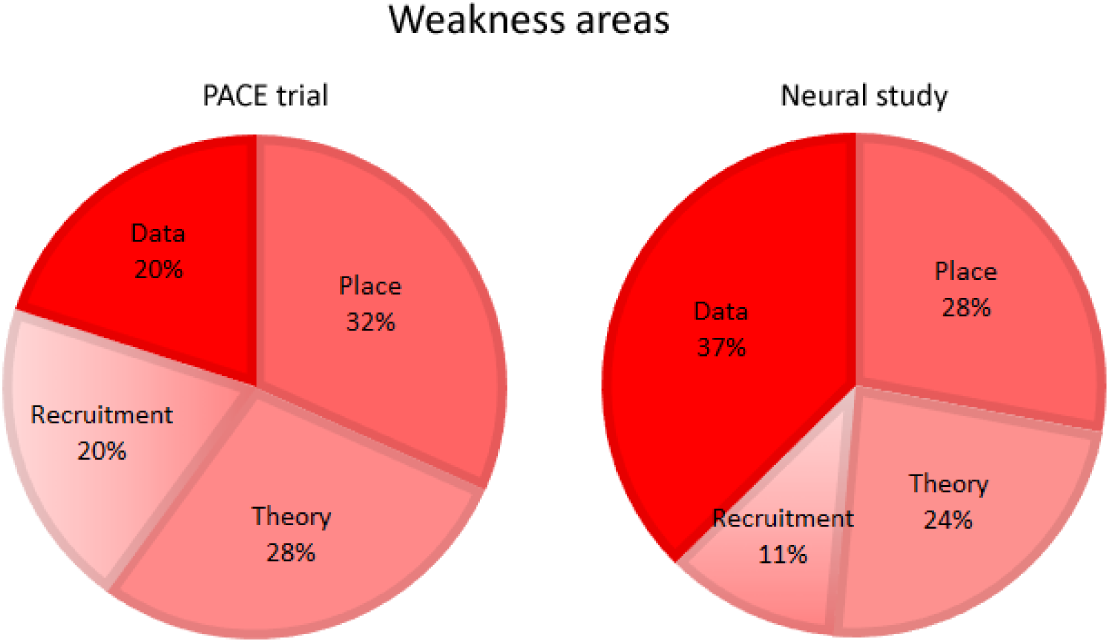
The proportion of weakness areas in PACE and NS.

### 4.6. Grants at risk of bias

The proportion of ME/CFS grants declared in psychosocial publications were at 56% (*n* = 722; number of articles; search dates 1951 to 25 March 2017) and 44% (*n* = 568) for cellular. The proportion of grants biased in presenting evidence for psychosocial were at 71.8% and 28.2% for cellular (OR = 2.16; fig. 7). Grants biased in presenting justifications were at 86.4% for psychosocial and 13.6% for cellular (OR = 4.76; fig. 7).

**Figure 7:**
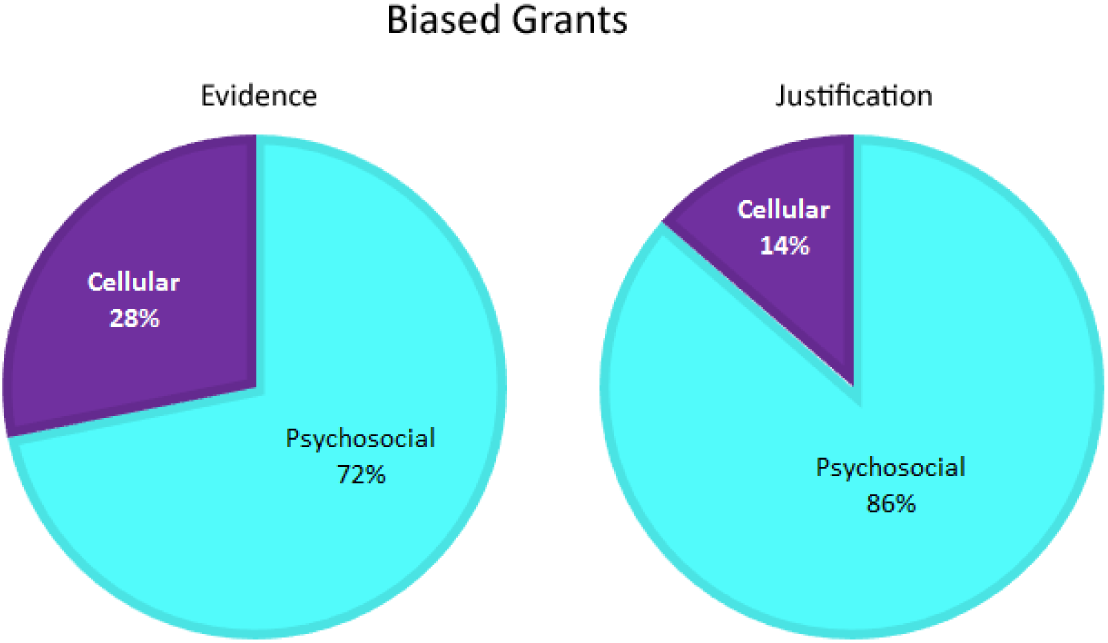
The proportion of bias in ME/CSF grants between research areas: psychosocial and cellular.

## 5. Discussion

The results suggest psychosocial ME/CFS trials are more likely to engage in selective reporting than cellular ME/CFS trials. It confirms the concerns raised by ME/CFS groups that psychosocial interventions are harmful, and present questionable therapeutic benefits no different to a placebo (Childs *et. al.*, 2015; Lian & Nettleton, 2015). However, the results also suggest, cellular trials are also likely to engage in selective reporting, but its therapeutic benefit is difficult to assess, since no study as of yet have proposed a therapeutic agent (eg. drug) exclusively designed and marketed for treating ME/CFS (Collatz *et. al.*, 2016). Brurberg and colleagues (2014) propose the need for consistency in ME/CFS research by applying a diagnostic criteria, subject to a systematic evaluation. This need to adequately define ME/CFS is a recurring consensus among researchers (Jason, Boulton & Friedberg, 2010; Nacul *et. al.*, 2011; Johnston *et. al.*, 2013). Some propose a re-evaluation of domains and criteria in existing patient reported outcome measurements (PROMs) by considering subgroups to account for heterogeneity (different populations) in comorbid conditions (eg. thyroid issues) and patient characteristics (eg. children) (Nacul *et., al.,* 2011; Haywood, Staniszewska & Chapman, 2012; Johnston *et. al.*, 2014; Haywood, Collin & Crawley, 2014; Hvidberg *et. al.*, 2015; Murdock *et. al.*, 2016). Others propose the need to investigate biomarkers and immune-mediated networks in developing a prophylactic agent (Fuite, Vernon & Broderick, 2008; Schlauch *et. al.*, 2016; Vega *et. al.*, 2017; Armstrong *et. al.*, 2017).

If in the latter, cellular trials (eg. biomarkers, immune checkpoints etc.) may benefit from designing outcomes which are sensitive and specific for clinical practice, also safe and reproducible across clinical practice. If this is not feasible, then begin with pre-clinical models (eg. animal models) and confirm risk thresholds (endpoints; safety) to deter heterogeneity (diverse or novel methods) in clinical phases from biasing the true effect. On the other hand, psychosocial studies (itemising and validating PROMs), may benefit from ensuring operational aspects (place) are well documented and archived. This ensures selective reporting in measurement-systematic errors and misclassification effects (bias towards or away from the null) can be corrected, and do not misconstrue the true effect.

Selective reporting is a problem in research waste, and a bias appraisal on evidence and justifications is one way to bring light of this. Future studies may look at biases (reoccurring problems) across multiple studies consistent in each research area, so that oversight bodies (eg. grant committees) do not restrict researchers from freely conducting research by enforcing a general standard across all research areas to address research waste.

## Acknowledgements

I dedicate this to ME Awareness Week 2017. I would like to thank ME groups for your passionate advocacy and for sharing your stories with me. It’s truly inspiring. Thank you. A special shout out to @postersandme @johnthejack @TweetTipsforME @DrSpeedyandME @MyalgicE @ Lucibee @consent patient @chrisbrownofca1 @FrancieSaidSo @EllieArnott @JaneEBSmith @velogubbed; @MECFSquestions, @MEMilitant1 and @emmajoy6 for very helpful comments to this manuscript; and @DrFulli for attracting me to this issue.

## Supplementary material

Link to figures and codes: http://www.openwetware.org/wiki/User:Sonia

Lee/Notebook/ResearchWasteMECFS Link to slides: https://figshare.com/articles/PACE Trial Critical Appraisal slides /4685074

